# Large scale, low coverage population genomics approach to assess genetic variation and structure across vulnerable Swedish sand lizard populations

**DOI:** 10.1101/2025.09.12.675594

**Authors:** Mette Lillie, Patrik Rödin Mörch, Erik Wapstra, Sven-Åke Berglind, Jacob Höglund, Mats Olsson

## Abstract

Genomics has emerged as a powerful tool in conservation biology, offering valuable insights into the genetic diversity, population connectivity and adaptive potential of threatened species. Here, we apply a large-scale, low-coverage whole genome sequencing approach to the conservation genomics of the Swedish sand lizard (Lacerta agilis). Sand lizards occur in fragmented populations across Sweden, with varying population sizes and degrees of isolation, and face significant threats from habitat degradation and land use change. In this study, we sequenced 139 individuals to investigate genetic diversity, population structure and genetic load. We reveal strong population structure among Swedish sand lizards and identify populations with low genetic diversity and realised high genetic load. Genetic diversity in populations declined with increasing latitude, likely reflecting the historical northward expansion process and the subsequent population fragmentation and isolation. We observe strong population structure and reveal genetic distinctiveness of populations; findings with significant conservation implications. For example, the Sörmon population, a relict population in central Sweden, appears genetically distinct from all other populations. Further, the northern-most populations in Dalarna appear relatively small, with high genetic load, and thus may require intensive management via e.g., restored open successional habitat to aid population recovery. Our study adds to the growing field of conservation genomics and demonstrates the value of high-resolution whole genome approaches to the conservation efforts for threatened populations. Finally, our research contributes crucial insights for the conservation management of this vulnerable species in Sweden.

## Introduction

The progression of conservation genomics from conservation genetics has been underway for over a decade now, driven by the rapid development in sequencing technologies and bioinformatic methods (Primmer 2009; Allendorf, et al. 2010; McMahon, et al. 2014; Hohenlohe, et al. 2021). Genomic approaches enable comprehensive estimation and comparison of genetic diversity and differentiation across populations, helping to identify those at risk due to reduced standing genetic variation and diminished adaptive potential.

Genomics can also resolve fine-scale population structure, identifying genetically diverse and distinct populations that may require tailored management strategies to ensure their conservation. Conservation genomics applications are exemplified by recent studies resolving population structure, migration and admixture (Yang, et al. 2022; Gose, et al. 2024), historical inbreeding and genetic load (Smeds, et al. 2024), uncovering ancestry and adaptive loci (van der Valk, et al. 2024), as well as identifying genetic associations to infectious diseases that threaten wild populations (Batley, et al. 2021). Whole genome sequencing offers an unprecedented marker density and surveys a wide diversity of genetic variations not limited to single nucleotide polymorphisms, increasing their power for the detection of signatures of selection and local adaptation as well as for the identification of the genetic basis of phenotypic traits and diseases (Fuentes-Pardo and Ruzzante 2017). In the face of the ongoing biodiversity crisis and the growing number of populations requiring conservation interventions, incorporating modern genomic technologies offers significant benefits to conservation efforts. Here, we apply a low-coverage conservation genomics approach to the vulnerable populations of sand lizards (*Lacerta agilis*) in Sweden.

Sand lizards are an early successional species, found across a wide area of Eurasia. Swedish populations represent the northern extremity of their European distribution, founded approximately 10 000 years BP, when the climatic warming in northern Europe made immigration possible via a land bridge connecting Scandinavia to Europe (Björck 1995). This migration corridor was available until ca. 9 000 years BP, when the land bridge was submerged from glacial ice melt, forming the Baltic Sea (Björck 1995) and isolating the Swedish sand lizard populations from the European mainland. In the warm post-glacial period (between 8 000 and 2 500 years BP), favourable conditions facilitated the dispersal of the sand lizards throughout southern Sweden. Subsequent climate cooling in combination with the loss of open, sandy habitats resulted in population fragmentation and local extinctions (Gullberg, et al. 1998; Berglind 2004a). Populations in the south-east of Sweden are generally more continuously distributed, where climatic conditions and environmental features are more favourable, whereas in central Sweden, a small number of populations persist on remnant glaciofluvial sand deposits within pine heath forests (Berglind 2004a; Berglind, et al. 2015).

As early successional species, sand lizards in Sweden are predominately threatened by land-use change and forest management practices, which result in the loss of suitable open habitat patches for foraging and reproduction, leading to habitat fragmentation and population isolation. For species dependent on early-successional landscapes, such as the sand lizard, the availability of suitable open habitat patches depends on the frequency and the extent of environmental disturbances and their persistence (White and Pickett 1985). Natural disturbances such as fires and floods are key ecological drivers that generate such patches and thus the species that inhabit these dynamic landscapes are continuously affected by a cycle of disturbance and forest regeneration (Brawn, et al. 2001; Berglind 2004a). Fire suppression and afforestation in recent years has seen an increase in forest cover in Europe, which has negatively impacted open-habitat species, from plants, to insects and animals (Plieninger, et al. 2013; Melero, et al. 2016; Regos, et al. 2016; Palmero-Iniesta, et al. 2020).

The Swedish sand lizard is currently red-listed as a vulnerable species and managed under a national species conservation action plan (Berglind, et al. 2015). Early genetic analyses have shown that the Swedish sand lizard populations have lower genetic diversity at both neutral (microsatellite markers) and adaptive genetic markers, compared with Central European populations (Gullberg, et al. 1998; Madsen, et al. 2000). Here, we use the high-quality sand lizard reference genome and low-coverage, individual-based whole-genome sequencing of 139 individuals and from two sampling time points to investigate genetic diversity, genetic load and population structure among sand lizard populations from across their Swedish distribution (Figure 1; Table 1).

**Figure 1.**
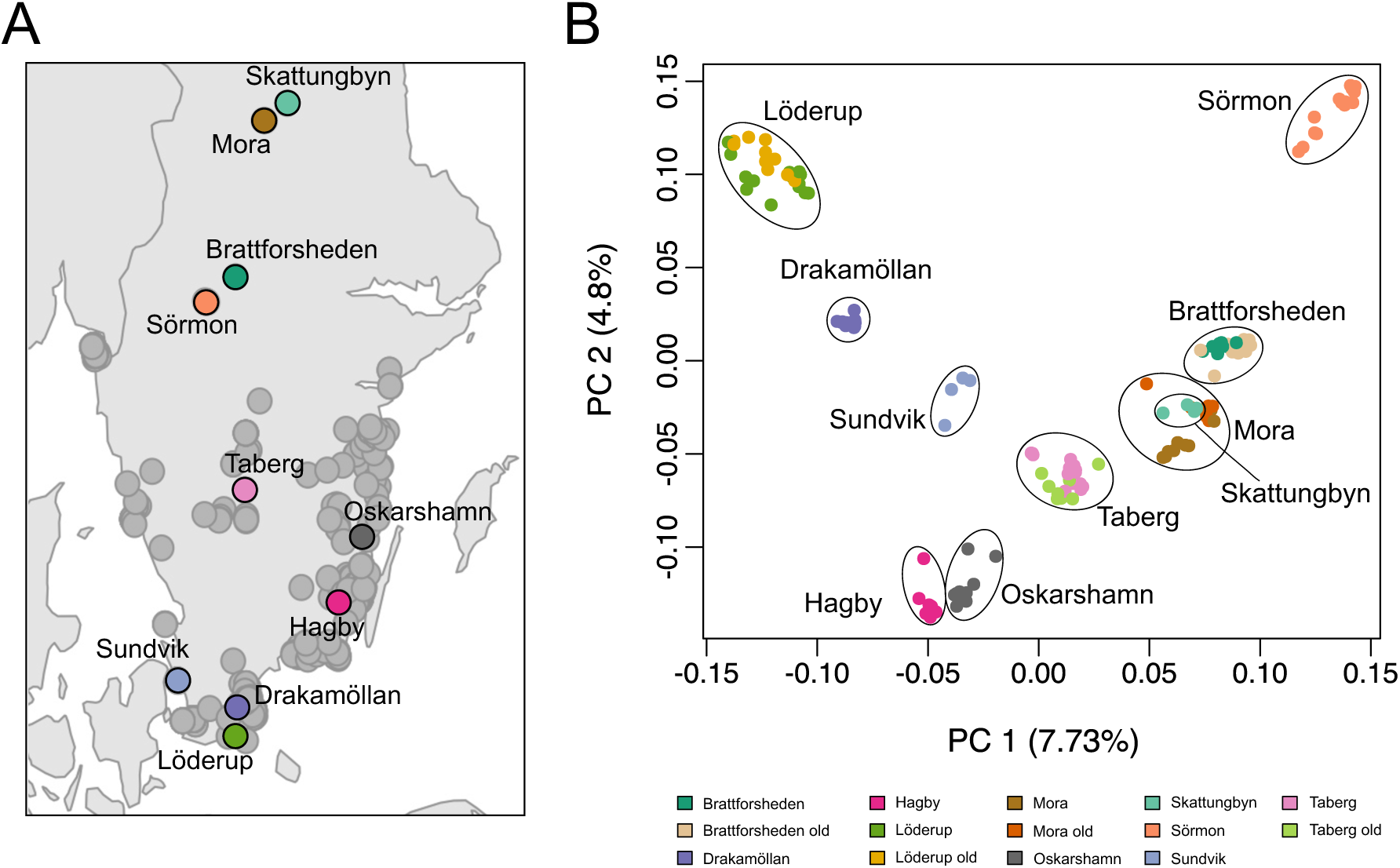
**A.** Map of southern Sweden showing sampling locations included in this study. **B.** Principal Component Analysis of whole genome sequencing data for PC1 and PC2. Percentage of variance explained by each PC given in parentheses.

**Table 1.**
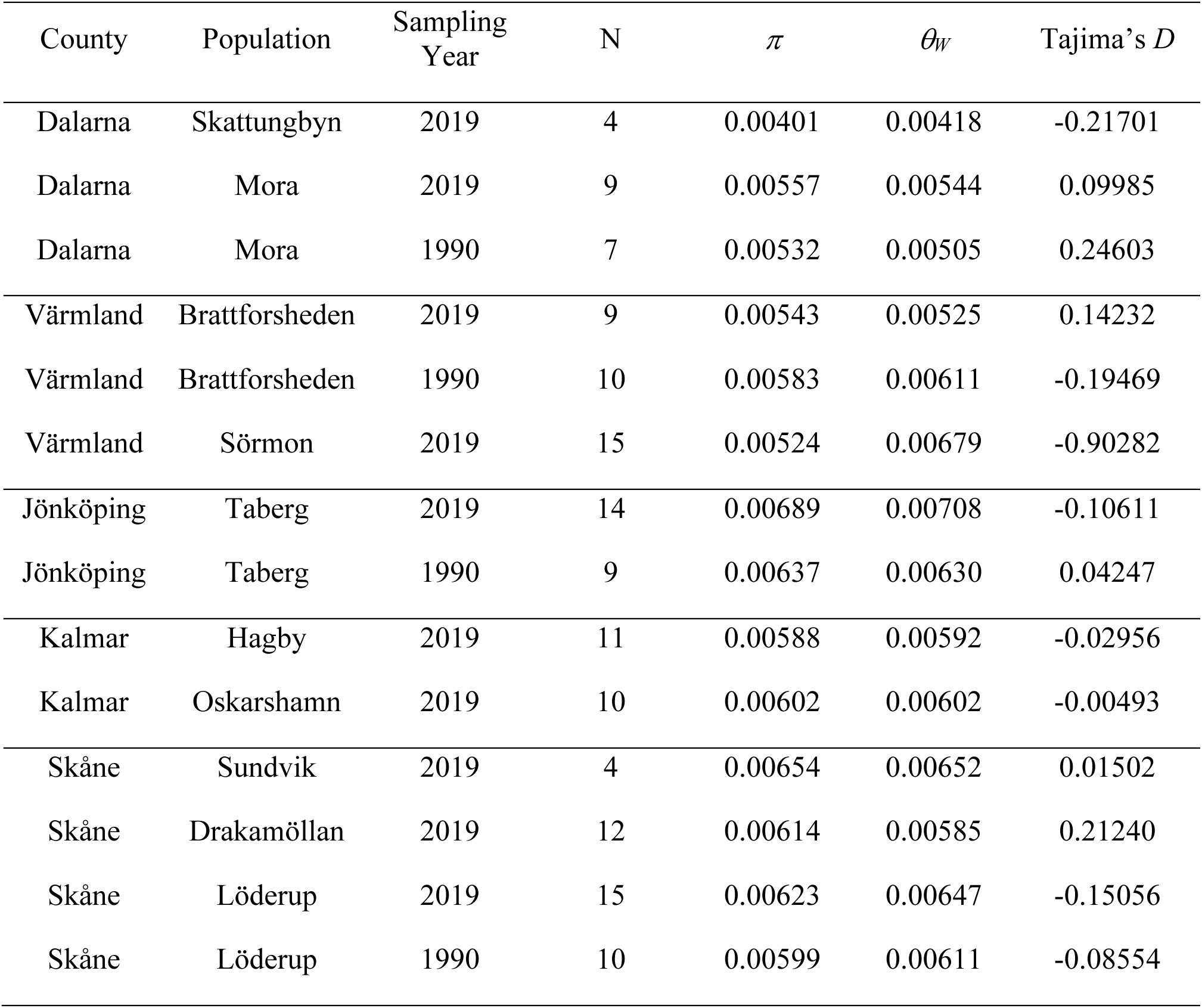
Population information with diversity statistics, including Swedish county, population name, sampling year, number of sampled individual (N), estimates of genetic diversity (nucleotide diversity, *π*; Watterson’s *8*, *8_W_*) and Tajima’s *D*.

## Material and methods

### Samples

Four Swedish populations had been previously sampled in the early 1990s (Gullberg, et al. 1998), totalling 36 samples, including Dalarna (Mora), Värmland (Brattforsheden), Jönköping (Taberg), Skåne (Löderup) (Figure 1; Table 1). During 2019, these populations were sampled again, as well as additional populations in Skåne (Drakamöllan; Sundvik), Kalmar (Oskarshamn; Hagby), Värmland (Sörmon) and Dalarna (Skattungbyn). In total, 103 samples were collected from 10 modern sand lizard populations (Figure 1; Table 1; Supplementary table A1). Tail tissue from a sand lizard sample from Bulgaria collected as part of a previous study (Henle, et al. 2017) was also used in this study.

### Whole genome sequencing

Samples from pre-2000 had been extracted using a salt-chloroform extraction as previously described (Gullberg, et al. 1998). Genomic DNA from the remaining samples was extracted using the QIAGEN DNeasy Blood and Tissue kit using the manufacturers protocol. DNA quality and concentration was assessed via NanoDrop 2000 Spectrophotometer (Thermo Scientific). Sequencing libraries were constructed from 10 ng of input DNA, using an in-house Tn5 protocol adapted from Picelli et al. (2014). This is a low-cost protocol for whole-genome sequencing library preparation, requiring only modest molecular experience.

Libraries were uniquely indexed (using combinatorial dual indexes) and pooled in equimolar multiplexes of up to 65 individuals (four pools in total; aiming for approximately 5 × per individual for the Swedish individuals, and 40 × for the European sample). Each pool was sequenced on an Illumina NovaSeq S2 lane at the SNP&SEQ Technology Platform (Uppsala, Sweden) and reads were separated bioinformatically based on their index.

### Bioinformatic analyses

Unmapped bams were generated for each sample and Illumina adaptors marked using picard v2.10.3 (http://broadinstitute.github.io/picard/index.html). These were then mapped to the sand lizard reference genome (Genbank accession: GCA_009819535.1) using bwa mem -M v0.7.17 (Li and Durbin 2009). The sand lizard genome was assembled using PacBio, Illumina, Hi-C and Bionano technologies, encompasses 1.39 Gbp of assembled genome, including 18 chromosomes, the Z and W sex chromosomes, mitochondrial genome, and eight unplaced shorter scaffolds (over 99.9% of the genome is assigned to chromosomes), with full gene annotation. This high-quality, high-resolution genome provides an invaluable asset for sand lizard population genomics. After mapping, picard was used to sort and mark duplicates in the mapped bam files. Samtools v1.18 depth (Danecek, et al. 2021) was used to compute coverage per sample. Bedtools v2.26.0 genomecov (Quinlan and Hall 2010) was used to compute sites covered by at least 1, 2, 3, 4, 5, 10 or 20 reads. The repeatmasker output from the sand lizard reference genome release was used to filter out repeat regions/low complexity regions from analyses.

### Nucleotide diversity

ANGSD v0.933 (Korneliussen, et al. 2014) was used to estimate genotype likelihoods from our low-coverage sequencing data, using the GATK algorithm (-gl 2). We estimated theta for each population using the European sample as ancestral. The ancestral state was set using ANGSD (options: -dofasta 2 -setMinDepth 20 -setMaxDepth 150 -minMapQ 20 -minQ 20 - remove_bads 1 -uniqueOnly 1). Allele frequencies per population was estimated using ANGSD options -dosaf 1 -GL 2 -minMapQ 20 -minQ 20 -remove_bads 1 -uniqueOnly 1 - dumpCounts 2 -doMajorMinor 5 -doMaf 2. We then used ANGSD realSFS to generate the unfolded site frequency spectrum per population, which could be used with the saf2theta and do_stat command to output diversity statistics for each chromosome. Diversity estimates (Wattersons *8, 8_W_*; nucleotide diversity, π) for each chromosome was divided by the number of sites per window to recover an unbiased estimate of diversity.

### Genetic load

To estimate the genetic load in each population, genotypes were called using ANGSD for each individual (-gl 2 -doPost 1 -doMajorMinor 1 -doMaf 1 -dobcf 1 --ignore-RG 0 -doGeno 1 -doCounts 1 -snp_pval 1e-6 -minMapQ 30 -minQ 20). The mutational effect for each called variant was estimated using SnpEff v5.2a (Cingolani, Platts, et al. 2012) using a local database built on the sequence and annotation of the sand lizard genome. Variant positions annotated with low, modifier, moderate or high impact were extracted using SnpSift v5.2 (Cingolani, Patel, et al. 2012), and the number of heterozygous (masked genetic load) and homozygous (realised genetic load) variants for each impact class and individual was summed per population. Genetic load proportion was calculated by dividing by the number of sites called within each population.

### Population structure and differentiation

Population structure was investigated using PCAngsd v1.11 (Meisner and Albrechtsen 2018; Meisner, et al. 2021). Genotype likelihoods were generated in ANGSD (-doGlf 2 -GL 2 - doMajorMinor 4 -doMaf 1 -uniqueOnly 1 -remove_bads 1) for sites with minor allele frequency >= 0.05 (supplied as a sites file). Principal component analysis (PCA) was then conducted in PCAngsd. Genotype likelihoods were also used for admixture analysis using NGSadmix v32 (Skotte, et al. 2013) for *K* values from 2 to 13. Best *K* was estimated using delta*K* from resulting likelihood values (Evanno, et al. 2005). Pairwise global *F*_ST_ estimates and window-based *F*_ST_ scans (100 kb windows with 50 kb step size) were calculated in ANGSD from the allele frequencies per population (generated for nucleotide diversity statistics, see above), using realSFS fst stats and fst stats2. Pairwise comparisons were focused on comparisons between modern populations, and within-population temporal comparisons. Z-transformed *F*_ST_ values were calculated and z*F*_ST_ values above 5 were used to define candidate genomic regions that have experienced directional selection. Genes within putative selected regions were identified and exons extracted from the reference genome annotation to generate a -sites file in order to estimate allele frequency differences within exons using angsd -gl 2 -doMajorMinor 4 -doMaf 2. Genes with an allele frequency difference greater than 0.6 between temporal sampling were identified and gene ontology terms retrieved from the PANTHER database (https://pantherdb.org/; accessed 2025/02/17).

## Results

Sequencing coverage for the final dataset was on average 6.10 × (median 6.14 ×) per individual, with over 128 samples (91%) having at least 5 × coverage (Supplementary Figure A1).

### Genetic diversity

Nucleotide diversity estimates (π) ranged from 0.00401 (Skattungbyn) to 0.00689 (Taberg), with generally less diversity in the northern populations, seen as a negative relationship between π and latitude (adjusted *R^2^*=0.5093, *p-*value=0.0123; Table 1; Figure 2A). A similar non-significant negative trend was seen at *8_W_* (adjusted *R^2^*=0.2681, *p-*value=0.0719). Values for Tajima’s D across populations all were close to 0 (between -0.22 and 0.25), with the exception of Sörmon, which had a slightly negative value of -0.902 (Table 1).

**Figure 2.**
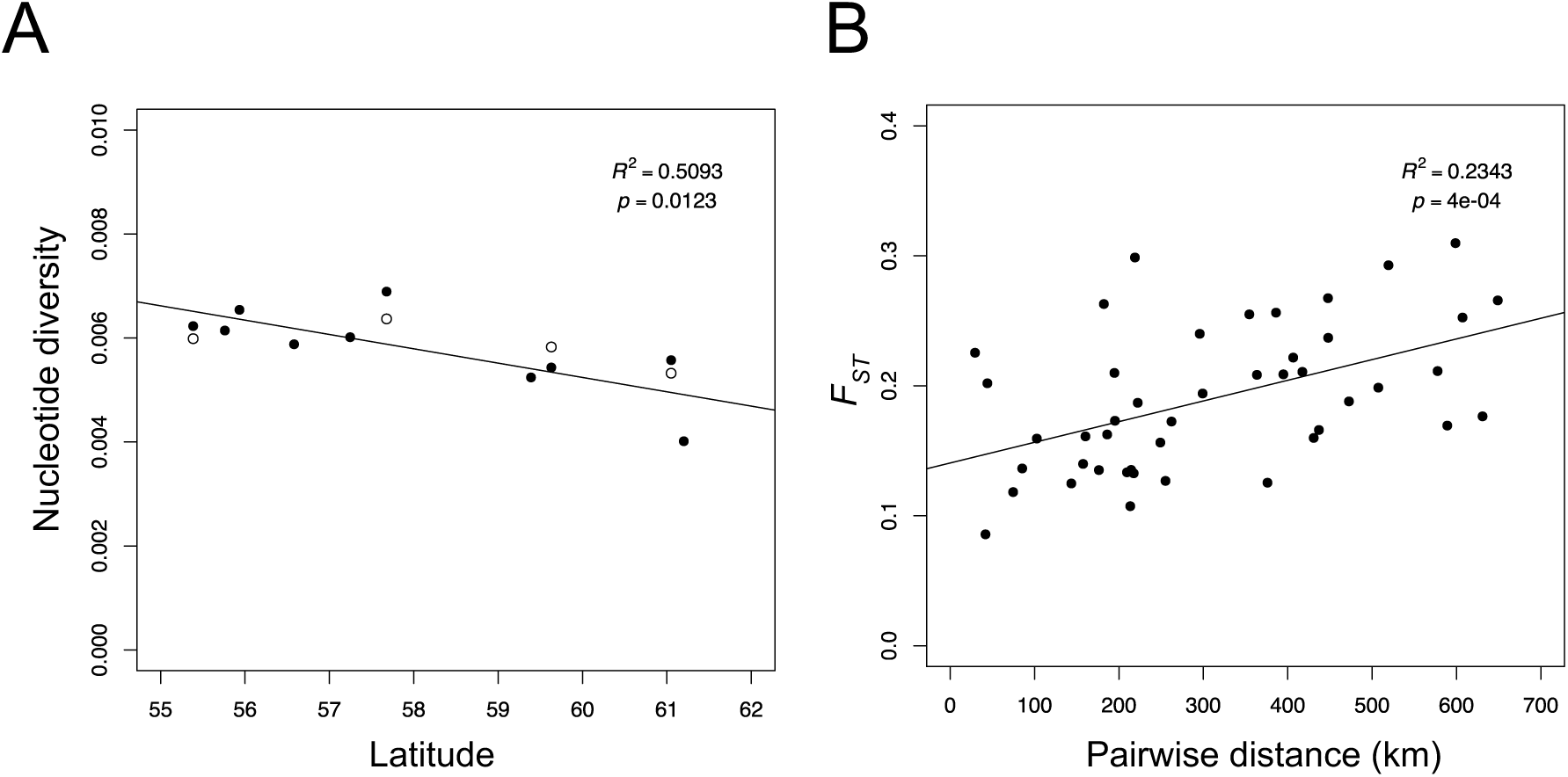
A. Nucleotide diversity (π) across latitudes of sand lizard populations for sampling in 2019 (filled circles) and 1990 (open circles). Linear regression line for the populations sampled in 2019 plotted, with *R*^2^ and *p*-value given. **B.** Genetic differentiation (*F_ST_*) versus geographic distance (haversine) between sand lizard population pairs sampled in 2019. Linear regression line plotted, with *R*^2^ and *p*-value given.

### Genetic load

Genetic load proportions were comparable across populations for both masked and realised load and between temporal samples within populations (Figure 3A; Supplementary Figure A2; Supplementary Figure A3). Masked and realised genetic load proportion was elevated in Skattungbyn for all impact classes.

**Figure 3.**
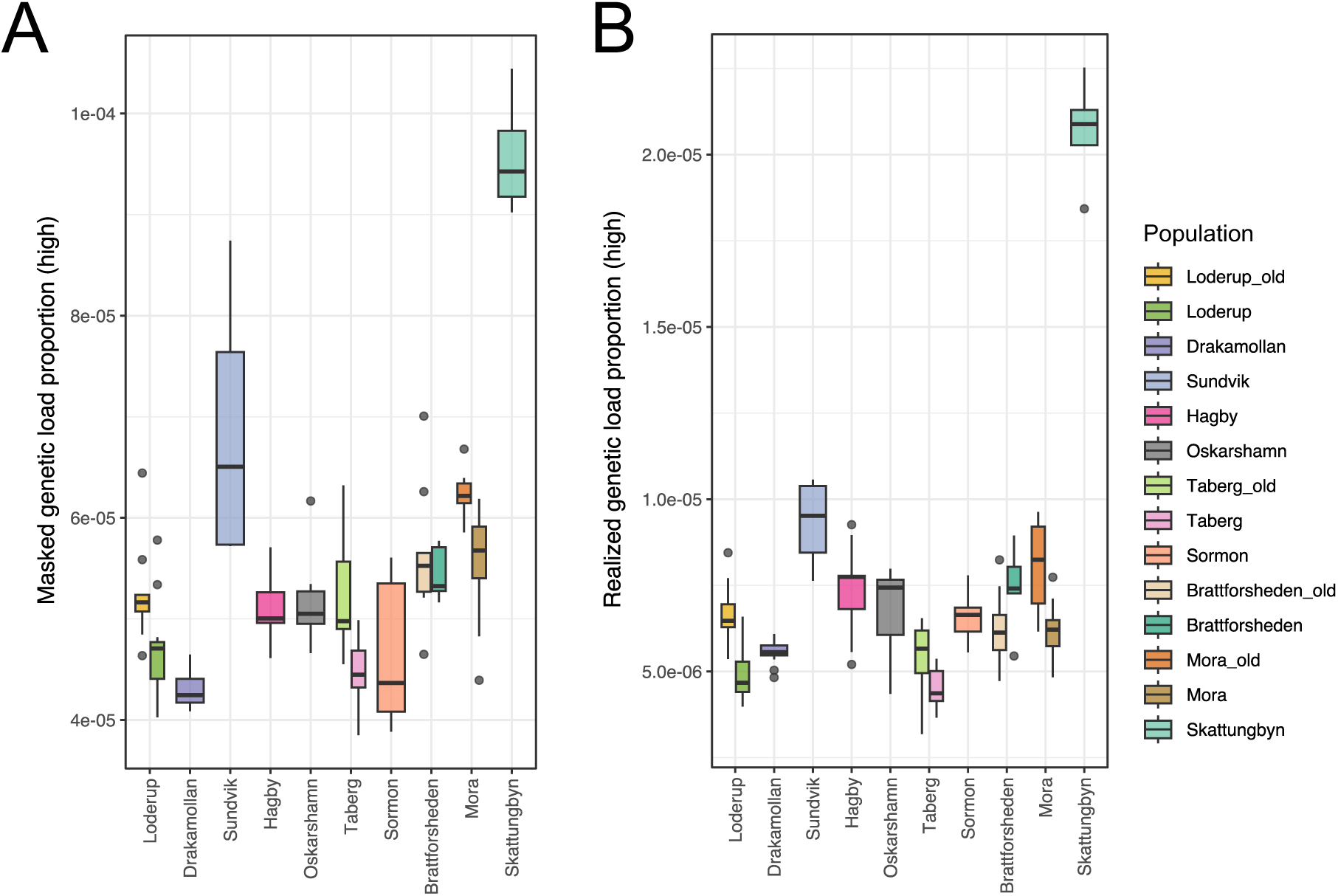
Genetic load proportions across Swedish sand lizard populations. **A.** Mean masked genetic load proportion (high impact deleterious mutations in heterozygous state) per population **B.** Realised genetic load proportion (high impact deleterious mutations in homozyogous state) per population. Populations ordered by increasing latitude.

### Genetic differentiation

Swedish sand lizard populations showed strong genetic structure in the PCA (Figure 1B). The PCA plot of PC1 and PC2 (Figure 1B) shows that individuals cluster by population, with the exception of Skattungbyn and Mora, which clustered together (both populations are in Dalarna, approx. 30 km apart). Temporal samples (samples collected 1990 and 2019) clustered by population. Optimal number of genetic groups (*K*) in admixture analysis was four, having the highest value of delta*K* (Supplementary Figure A4; Supplementary Figure A5). At *K*=4, admixture plots showed a general genetic divide between southern, eastern, and northern populations, with the Sörmon samples forming a distinct genetic cluster (Supplementary Figure A4). As such, Löderup appeared “southern”, Hagby and Oskarshamn appeared “eastern” and Brattforsheden, Mora and Skattungbyn appeared “northern”.

Drakamöllan and Taberg appeared admixed, such that Drakamöllan admixture proportions included both “southern” and “eastern”, while Taberg admixture proportions were predominately “northern” and “eastern”.

Population differentiation (global weighted *F_ST_*) between population pairs varied from 0.0904 (Drakamöllan-Löderup) to 0.3097 (Sundvik-Skattungbyn) (Table A2). Pairwise differentiation showed correlation with geographic distance between population pairs, likely reflecting isolation by distance (*R^2^*=0.2503, *p-*value<0.001 Figure 2B; Supplementary table A2). Values for *F_ST_* were relatively high in the northern populations, for example, *F_ST_* was 0.2254 between the populations Skattungbyn and Mora (Dalarna), which are located approximately 30 kms apart, and 0.2020 between the populations Brattforsheden and Sörmon (Värmland), which are only 45 kms apart (Table A2). In contrast, differentiation among population-pairs from southern Sweden was lower, e.g., *F_ST_* was 0.1182 between the Kalmar populations Hagby and Oskarshamn (approximately 85 km apart).

Levels of differentiation between temporal sampling within populations was comparably lower, with global genome-wide *F_ST_* values varying between 0.0210 and 0.0447 (Table 2). Window-based *F_ST_* scans of the genome revealed several peaks of differentiation within populations over time (Supplementary Figure A6), potentially indicating regions of genome under directional selection. A total of 18 genes within *F_ST_* peaks were identified with pronounced allele frequency differences between temporal samples in Mora, 33 in Brattforsheden, 12 in Taberg and 11 in Löderup. Genes were associated with diverse functions, including development, cellular regulation, neuronal signal and development, metabolism and immune response (Supplementary table A3).

**Table 2.** Global *F_ST_* (weighed) between temporal sampling of sand lizard populations, sampled in 1990 and 2019.

| Population | $F_{ST}$ |
| --- | --- |
| Mora | 0.0372 |
| Brattforsheden | 0.0390 |
| Taberg | 0.0447 |
| Löderup | 0.0210 |

## Discussion

Conservation genomics applies advanced DNA sequencing approaches to explore the complex genetic landscapes of endangered species, providing valuable insights that can inform and guide conservation management actions. In this study, we apply low-coverage whole genome sequencing to the vulnerable populations of sand lizard in Sweden. We reveal strong population structure among Swedish sand lizards and identify populations with low genetic diversity and high realised genetic load. These findings have significant conservation implications for a species at the northern climatic edge of its distribution and red-listed in Sweden.

We observed a negative relationship between genetic diversity (π and *8_W_*) with latitude in Swedish sand lizard populations, reflecting patterns observed previously in Swedish sand lizards using microsatellite markers (Gullberg, et al. 1998). This follows expectations that the expansion from central European refugial areas to Sweden was characterised by repeated founder effects, resulting in populations with lower diversity at the edge of a species’ range (Hewitt 2000; Eckert, et al. 2008). Swedish sand lizard populations were founded via a southern route with genetic diversity limited by the one migration period during a period of favourable conditions that facilitated northward expansion. The higher latitude populations in Värmland and Dalarna, thus, contained the lowest genetic diversity, similar to patterns observed in other vertebrates (Yiming, et al. 2021; Fonseca, et al. 2023). These populations in Värmland and Dalarna have been described as isolated relicts, confined to remnant glaciofluvial sand deposits in pine heath forests that persisted when the climate cooled (Berglind 2004a). It should be noted that genetic diversity for Skattungbyn may be underestimated due to the low sample size. This population was particularly difficult to sample, however, despite concerted effort, suggesting small population size.

Genetic load proportions showed no significant difference across latitude. Thus, although genome-wide genetic diversity appears to decline with latitude as a result of the historical expansion process and subsequent fragmentation and isolation, these populations do not appear to have accumulated significantly more genetic load, with the exception of Skattungbyn. Estimates for masked and realised load were greatest in Sundvik and Skattungbyn, but these may be inflated due to the lower sample sizes. The high genetic load estimates in Skattungbyn (present as both realised and masked load) may reflect that this population contains relatively more deleterious variation, which may indicate serious conservation consequences. Realised load is likely to impact population-level fitness (van Oosterhout 2020; Kyriazis, et al. 2021). Persistent small population size and inbreeding will, in turn, convert the proportionally high standing masked load to more realised load, further exacerbating the situation in the current Skattungbyn population. Inbreeding in other sand lizard populations has been associated with increase malformations in hatchlings (Olsson, et al. 1996) and lower hatching success (Bererhi, et al. 2019). While our results provide valuable insights into the potential genetic risks, caution is warranted in their interpretation due to the reliance on low-coverage sequencing data. Higher coverage whole genome sequencing can be used to confirm these findings, and at the same time, allow for inbreeding to be quantitatively assessed via runs of homozygosity analysis. Although purely genomic analyses of genetic load are insufficient to fully evaluate viability (Kardos, et al. 2024), our results call for further work in the Skattungbyn population to e.g., investigate individual fitness, population demography and functional genetic diversity. Such additional data would strengthen our ability to estimate Skattungbyn’s vulnerability to extinction and inform more targeted conservation strategies. It should be noted that small-scale habitat restorations have been undertaken for this population for more than ten years, but these have not reached the recommendations outlined in the Swedish conservation action plan for the sand lizard. The plan recommends the creation of large (5-10 ha), high-quality habitats, which has proven successful in Brattforsheden to aid population recovery (Berglind, et al. 2015).

Swedish sand lizard populations show high genetic differentiation (*Fst*), with strong population structure evident from PCA and admixture analysis, reflecting the prolonged fragmentation of populations. Admixture analysis indicates a southern, eastern and northern genetic divide of populations (*K*=4), with admixture between these regions in e.g. Drakamöllan and Taberg. These patterns may reflect the original dispersal throughout Sweden and the differentiation of populations along different colonisation routes during the expansion process. Sand lizard populations likely went extinct except in the most favourable localities during the cooling of the Holocene with climate change and habitat fragmentation impacting dispersal and gene flow (Gullberg, et al. 1998).

Sörmon appears genetically distinct from all other populations, even Brattforsheden, which is separated by only 45 km. Genome-wide *F_ST_* estimates between Sörmon and Brattforsheden (0.2020) were greater than that between Mora and Brattforsheden (0.1611), which are over 150 kms apart. This may indicate that the Sörmon population was established from a separate colonization route to that of Brattforsheden and the populations of Dalarna, which has produced the deep divergence evident in the genetics of the modern-day populations.

Potentially, Sörmon was colonized via a western route, and sampling of the sand lizard populations on Swedeńs western and north-western coast would help confirm this theory. This finding also has implication to the conservation management of sand lizard populations. The Dalarna populations are small and require intensive restoration actions to generate large, open successional habitat of high quality in the sandy glaciofluvial areas, which have supported the recovery of population sizes in the Brattforsheden populations (Berglind, et al. 2015). Habitat restoration could also be combined with translocation of eggs or individuals (Berglind 2004b). Should conservation efforts consider genetic rescue of populations in Dalarna, Brattforsheden may be considered as donor population, given the apparent historical relatedness of the populations. Conversely, the apparent genetic distinctness of Sörmon may indicate a high conservation value of this population, which has undergone distinct local adaptation and drift processes and would preclude admixture with other populations of our study.

We observed high genetic diversity (π and *8_W_*) in Taberg, which is an isolated population located in the centre of the sand lizard distribution (Gullberg, et al. 1998). Admixture proportions in this population for *K*=4 indicate contributions from all genetic clusters, with predominately eastern and northern genetic admixture compositions. This suggests that the Taberg population has been shaped by secondary contact of different colonisation routes, increasing the diversity in this population (though care should be taken to overestimate admixture results, see Lawson, et al. (2018)). A favourable environment in Taberg, capable of sustaining a large sand lizard population size may also contribute to the persistence of genetic diversity. This population inhabits the site of a retired iron-ore mine, now a nature reserve since 1985. The remnants from mining activities and mining waste throughout the site facilitates the persistence of open habitat patches within surrounding forest. This environmental mosaic provides favourable microclimates for thermoregulation, foraging, mating behaviours, and protection (Nemes, et al. 2006) and maintain a relatively large population size, retaining genetic diversity despite geographic isolation.

Temporal sampling in 1990 and 2019 allowed us to investigate genetic diversity and directional selection in four populations. Populations showed very little change in genetic diversity over time with low genome-wide differentiation. This suggests that populations have, in general, maintained genetic diversity through time, and may not have experienced drastic demographic effects and exacerbated genetic drift in recent decades. This also allowed us to investigate regions that appear under directional selection via window-based *F_ST_* estimation and outlier analysis. Genes within peaks across these populations had diverse roles including development, cellular signalling and regulation, and neuronal signalling and development. Within Löderup, three from 11 potentially selected genes were involved in the immune response, including the polymeric immunoglobulin receptor (PIGR) and complement receptors CR1 and CR2. PIGR plays an important role in mucosal immune protection and the maintenance of host mucosal barrier integrity, protecting hosts from pathogens while promoting a healthy relationship with commensal bacteria (Phalipon and Corthésy 2003). The complement receptors (CR) are critical regulators of the immune response, regulating leukocyte recruitment and migration, phagocytosis, and inflammation, playing roles in both the innate and adaptive immune response (Erdei, et al. 2009; Holers 2014; Vandendriessche, et al. 2021). Furthermore, polymorphism of CR1 and CR2 have been linked to susceptibility to infectious disease in humans (Luo, et al. 2019; Diep, et al. 2023).

The shifts in allele frequencies in these genes in Löderup could indicate a change in the pathogenic pressure in this population. Caution should be exercised interpreting these results, however, as *F_ST_-*based genome scans can be confounded by effect of drift in regions of low recombination (Booker, et al. 2020).

With the advance in sequencing technologies and decline in associated costs, genomics is poised to play an integral role in biodiversity conservation (Theissinger, et al. 2023). Large-scale genome assembly initiatives, such as Darwin Tree of Life (Consortium, et al. 2022), Earth Biogenome Project (Lewin, et al. 2018), the European Reference Genome Atlas (Formenti, et al. 2022) and the Vertebrate Genomes Project (Rhie, et al. 2021), are generating high quality reference genomes for hundreds of organisms, providing the valuable foundation for conservation genetics research programs (Brandies, et al. 2019). We developed a cost-effective approach that capitalises on the available high-quality sand lizard genome, employing in-house library preparation and low-coverage sequencing. This relatively low-cost approach may be highly compatible with the restricted budgets of conservation programs. This approach is highly scalable and flexible, and could be easily adapted for higher coverage sequencing, if required, for example, for more comprehensive inbreeding analyses via runs of homozygosity. In applying this method, we generated high-resolution, fine-scale genomic information with improved genetic insights compared to previous methods (Gullberg, et al. 1997; Gullberg, et al. 1998) with direct implications for the conservation management of sand lizards in Sweden. We identify populations requiring immediate conservation action and highlight genetically distinct populations that may warrant independent management strategies. This study demonstrates the utility of this approach to conservation genomics and we encourage its adoption by others in the field.

## Acknowledgements

We thank Elisabeth Bak-Olsson for assistance in the field (Taberg). We thank Ronny Wolf and Martin Schlegel (University of Leipzig, Leipzig, Germany) for sharing sand lizard samples from their earlier European study (Henle, et al. 2017). This study received financial support from Wilhelm & Martina Lundgrens vetenskapsfond, Stiftelsen för zoologisk forskning, Stiftelsen J A Wahlbergs minnesfond (through Kungliga Vetenskapsakademien, KVA), the Nilsson-Ehle Endowment, Adlerbertska forskningsstiftelsen (through Kungliga Vetenskaps-och Vitterhets-Samhället, KVVS) to ML. ML was supported by the Carl Tryggers foundation (CTS 16:343) and the Swedish Research Council (2021-04238). SÅB was supported by the Swedish Environmental Protection Agency. MO was supported by the Swedish Research Council (2021-04880). JH was supported by the Swedish Research Council (2023-05073) and Formas (2023-01050). EW was supported by the Australian Research Council.

Sequencing was performed by the SNP&SEQ Technology Platform in Uppsala. The facility is part of the National Genomics Infrastructure (NGI) Sweden and Science for Life Laboratory. The SNP&SEQ Platform is also supported by the Swedish Research Council and the Knut and Alice Wallenberg Foundation. Computations and data handling were enabled by resources in projects (NAISS 2024/5-209, NAISS 2023/22-419, NAISS 2024/6-81, NAISS 2023/23-604) provided by the National Academic Infrastructure for Supercomputing in Sweden (NAISS) at UPPMAX, funded by the Swedish Research Council through grant agreement no. 2022-06725.

## Data availability statement

raw sequencing data with associated metadata are available via the Sequence Read Archive (SRA, https://www.ncbi.nlm.nih.gov/sra/; Bioproject: PRJNA1275812).

## Ethics approval statement

This research was conducted at the University of Gothenburg under scientific research permit (Dnr 5.8.18-12538/2017) issued by the Animal Ethics Committee at the University of Gothenburg, Sweden. Permits to sample individuals from each population (Ansökan om dispens från fridlysningsbestämmelserna i 4-9 §§ artskyddsförordningen (2007:845)) were obtained from relevant Country Administrative Boards (Länsstyrelsen) (permit numbers: Dalarna: 522-13624-2018; Jönköping: 522-9603-2018; Kalmar: 522-9389-2018; Värmland: 522-942-2019; Västra Götaland: 522-42582-2018; Skåne: 522-35378-2018).

## Data Accessibility and Benefit-Sharing section

### Data Accessibility Statement

Raw sequencing data with associated metadata are available via the Sequence Read Archive (SRA, https://www.ncbi.nlm.nih.gov/sra/; Bioproject: PRJNA1275812).

### Benefit-Sharing Statement

Benefits Generated: Benefits from this research accrue from the sharing of our data and results on public databases as described above.

## References

Allendorf FW, Hohenlohe PA, Luikart G. 2010. Genomics and the future of conservation genetics. Nature Reviews Genetics 11:697–709.

Batley KC, Sandoval-Castillo J, Kemper CM, Zanardo N, Tomo I, Beheregaray LB, Möller LM. 2021. Whole genomes reveal multiple candidate genes and pathways involved in the immune response of dolphins to a highly infectious virus. Molecular Ecology 30:6434–6448.

Bererhi B, Wapstra E, Schwartz TS, Olsson M. 2019. Inconsistent inbreeding effects during lizard ontogeny. Conservation Genetics 20:865–874.

Berglind S-Å. 2004a. Area-sensitivity of the sand lizard and spider wasps in sandy pine heath forests - umbrella species for early successional biodiversity conservation? Ecological Bulletins 51:189–207.

Berglind S-Å. 2004b. Sand Lizard (Lacerta agilis) in Central Sweden: Modeling Juvenile Reintroduction and Spatial Management Strategies for Metapopulation Establishment. In: AkÇakaya HR, Burgman MA, Kindvall O, Wood CC, Sjögren-Gulve P, Hatfield JS, McCarthy MA, editors. Species Conservation and Management: Case Studies. New York: Oxford University Press. p. 326–339.

Berglind S-Å, Gullberg A, Olsson M. 2015. Conservation Action Plan for the Sand Lizard (Lacerta agilis) in Sweden (In Swedish with English summary). Naturvårdsverket, Rapport 6597.

Björck S. 1995. A review of the history of the Baltic Sea, 13.0-8.0 KA BP. Quaternary International 27:19–40.

Booker TR, Yeaman S, Whitlock MC. 2020. Variation in recombination rate affects detection of outliers in genome scans under neutrality. Molecular Ecology 29:4274–4279.

Brandies P, Peel E, Hogg CJ, Belov K. 2019. The Value of Reference Genomes in the Conservation of Threatened Species. Genes 10:17.

Brawn JD, Robinson SK, Thompson FR. 2001. The role of disturbance in the ecology and conservation of birds. Annual Review of Ecology and Systematics 32:251–276.

Cingolani P, Patel VM, Coon M, Nguyen T, Land SJ, Ruden DM, Lu X. 2012. Using Drosophila melanogaster as a Model for Genotoxic Chemical Mutational Studies with a New Program, SnpSift. Frontiers in Genetics 3:35.

Cingolani P, Platts A, Wang LL, Coon M, Nguyen T, Wang L, Land SJ, Lu XY, Ruden DM. 2012. A program for annotating and predicting the effects of single nucleotide polymorphisms, SnpEff: SNPs in the genome of Drosophila melanogaster strain w^1118^; iso-2; iso-3. Fly 6:80–92.

Consortium TDToLP, Blaxter M, Mieszkowska N, Di Palma F, Holland P, Durbin R, Richards T, Berriman M, Kersey P, Hollingsworth P, et al. 2022. Sequence locally, think globally: The Darwin Tree of Life Project. Proceedings of the National Academy of Sciences 119:e2115642118.

Danecek P, Bonfield JK, Liddle J, Marshall J, Ohan V, Pollard MO, Whitwham A, Keane T, McCarthy SA, Davies RM, et al. 2021. Twelve years of SAMtools and BCFtools. Gigascience 10.

Diep NT, Giang NT, Diu NTT, Nam NM, Khanh LV, Quang HV, Hang NT, Mao CV, Son HV, Hieu NL, et al. 2023. Complement receptor type 1 and 2 (CR1 and CR2) gene polymorphisms and plasma protein levels are associated with the Dengue disease severity. Scientific Reports 13:17377.

Eckert CG, Samis KE, Lougheed SC. 2008. Genetic variation across species’ geographical ranges: the central–marginal hypothesis and beyond. Molecular Ecology 17:1170–1188.

Erdei A, Isaák A, Török K, Sándor N, Kremlitzka M, Prechl J, Bajtay Z. 2009. Expression and role of CR1 and CR2 on B and T lymphocytes under physiological and autoimmune conditions. Molecular Immunology 46:2767–2773.

Evanno G, Regnaut S, Goudet J. 2005. Detecting the number of clusters of individuals using the software STRUCTURE: a simulation study. Molecular Ecology 14:2611–2620.

Fonseca EM, Pelletier TA, Decker SK, Parsons DJ, Carstens BC. 2023. Pleistocene glaciations caused the latitudinal gradient of within-species genetic diversity. Evolution Letters 7:331–338.

Formenti G, Theissinger K, Fernandes C, Bista I, Bombarely A, Bleidorn C, Ciofi C, Crottini A, Godoy JA, Höglund J, et al. 2022. The era of reference genomes in conservation genomics. Trends in Ecology & Evolution 37:197–202.

Fuentes-Pardo AP, Ruzzante DE. 2017. Whole-genome sequencing approaches for conservation biology: Advantages, limitations and practical recommendations. Molecular Ecology 26:5369–5406.

Gose MA, Humble E, Brownlow A, Wall D, Rogan E, Sigurosson GM, Kiszka JJ, Thostesen CB, Ijsseldijk LL, ten Doeschate M, et al. 2024. Population genomics of the white-beaked dolphin (Lagenorhynchus albirostris): Implications for conservation amid climate-driven range shifts. Heredity.

Gullberg A, Olsson M, Hegelstrom H. 1998. Colonization, genetic diversity, and evolution in the Swedish sand lizard, Lacerta agilis (Reptilia, Squamata). Biological Journal of the Linnean Society 65:257–277.

Gullberg A, Olsson M, Tegelstrom H. 1997. Male mating success, reproductive success and multiple paternity in a natural population of sand lizards: Behavioural and molecular genetics data. Molecular Ecology 6:105–112.

Henle K, Andres C, Bernhard D, Grimm A, Stoev P, Tzankov N, Schlegel M. 2017. Are species genetically more sensitive to habitat fragmentation on the periphery of their range compared to the core? A case study on the sand lizard (Lacerta agilis). Landscape Ecology 32:131–145.

Hewitt G. 2000. The genetic legacy of the Quaternary ice ages. Nature 405:907–913.

Hohenlohe PA, Funk WC, Rajora OP. 2021. Population genomics for wildlife conservation and management. Molecular Ecology 30:62–82.

Holers VM. 2014. Complement and Its Receptors: New Insights into Human Disease. Annual Review of Immunology 32:433–459.

Kardos M, Keller LF, Funk WC. 2024. What Can Genome Sequence Data Reveal About Population Viability? Molecular Ecology n/a:e17608.

Korneliussen TS, Albrechtsen A, Nielsen R. 2014. ANGSD: Analysis of Next Generation Sequencing Data. Bmc Bioinformatics 15.

Kyriazis CC, Wayne RK, Lohmueller KE. 2021. Strongly deleterious mutations are a primary determinant of extinction risk due to inbreeding depression. Evolution Letters 5:33–47.

Lawson DJ, van Dorp L, Falush D. 2018. A tutorial on how not to over-interpret STRUCTURE and ADMIXTURE bar plots. Nature Communications 9:3258.

Lewin HA, Robinson GE, Kress WJ, Baker WJ, Coddington J, Crandall KA, Durbin R, Edwards SV, Forest F, Gilbert MTP, et al. 2018. Earth BioGenome Project: Sequencing life for the future of life. Proceedings of the National Academy of Sciences 115:4325–4333.

Li H, Durbin R. 2009. Fast and accurate short read alignment with Burrows-Wheeler transform. Bioinformatics 25:1754–1760.

Luo J, Chen S, Wang J, Ou S, Zhang W, Liu Y, Qin Z, Xu J, Lu Q, Mo C, et al. 2019. Genetic polymorphisms in complement receptor 1 gene and its association with HBV-related liver disease: A case-control study. Gene 688:107–118.

Madsen T, Olsson M, Wittzell H, Stille B, Gullberg A, Shine R, Andersson S, Tegelstrom H. 2000. Population size and genetic diversity in sand lizards (Lacerta agilis) and adders (Vipera berus). Biological Conservation 94:257–262.

McMahon BJ, Teeling EC, Höglund J. 2014. How and why should we implement genomics into conservation? Evolutionary Applications 7:999–1007.

Meisner J, Albrechtsen A. 2018. Inferring Population Structure and Admixture Proportions in Low-Depth NGS Data. Genetics 210:719–731.

Meisner J, Albrechtsen A, Hanghøj K. 2021. Detecting selection in low-coverage high-throughput sequencing data using principal component analysis. Bmc Bioinformatics 22:470.

Melero Y, Stefanescu C, Pino J. 2016. General declines in Mediterranean butterflies over the last two decades are modulated by species traits. Biological Conservation 201:336–342.

Nemes S, Vogrin M, Hartel T, Öllerer K. 2006. Habitat selection at the sand lizard (Lacerta agilis): ontogenetic shifts. North-Western Journal of Zoology.

Olsson M, Gullberg A, Tegelstrom H. 1996. Malformed offspring, sibling matings, and selection against inbreeding in the sand lizard (Lacerta agilis). Journal of Evolutionary Biology 9:229–242.

Palmero-Iniesta M, Espelta JM, Gordillo J, Pino J. 2020. Changes in forest landscape patterns resulting from recent afforestation in Europe (1990–2012): defragmentation of pre-existing forest versus new patch proliferation. Annals of Forest Science 77:43.

Phalipon A, Corthésy B. 2003. Novel functions of the polymeric Ig receptor: well beyond transport of immunoglobulins. Trends in Immunology 24:55–58.

Picelli S, Bjorklund AK, Reinius B, Sagasser S, Winberg G, Sandberg R. 2014. Tn5 transposase and tagmentation procedures for massively scaled sequencing projects. Genome Research 24:2033–2040.

Plieninger T, Gaertner M, Hui C, Huntsinger L. 2013. Does land abandonment decrease species richness and abundance of plants and animals in Mediterranean pastures, arable lands and permanent croplands? Environmental Evidence 2:3.

Primmer CR. 2009. From Conservation Genetics to Conservation Genomics. In: Ostfeld RS, Schlesinger WH, editors. Year in Ecology and Conservation Biology 2009. p. 357–368.

Quinlan AR, Hall IM. 2010. BEDTools: a flexible suite of utilities for comparing genomic features. Bioinformatics 26:841–842.

Regos A, Domínguez J, Gil-Tena A, Brotons L, Ninyerola M, Pons X. 2016. Rural abandoned landscapes and bird assemblages: winners and losers in the rewilding of a marginal mountain area (NW Spain). Regional Environmental Change 16:199–211.

Rhie A, McCarthy SA, Fedrigo O, Damas J, Formenti G, Koren S, Uliano-Silva M, Chow W, Fungtammasan A, Kim J, et al. 2021. Towards complete and error-free genome assemblies of all vertebrate species. Nature 592:737-+.

Skotte L, Korneliussen TS, Albrechtsen A. 2013. Estimating Individual Admixture Proportions from Next Generation Sequencing Data. Genetics 195:693–702.

Smeds L, Huson LSA, Ellegren H. 2024. Structural genomic variation in the inbred Scandinavian wolf population contributes to the realized genetic load but is positively affected by immigration. Evolutionary Applications 17.

Theissinger K, Fernandes C, Formenti G, Bista I, Berg PR, Bleidorn C, Bombarely A, Crottini A, Gallo GR, Godoy JA, et al. 2023. How genomics can help biodiversity conservation. Trends in Genetics 39:545–559.

van der Valk T, Jensen A, Caillaud D, Guschanski K. 2024. Comparative genomic analyses provide new insights into evolutionary history and conservation genomics of gorillas. Bmc Ecology and Evolution 24.

van Oosterhout C. 2020. Mutation load is the spectre of species conservation. Nature Ecology & Evolution 4:1004–1006.

Vandendriessche S, Cambier S, Proost P, Marques PE. 2021. Complement Receptors and Their Role in Leukocyte Recruitment and Phagocytosis. Frontiers in Cell and Developmental Biology 9.

White PS, Pickett STA. 1985. Chapter 1 - Natural Disturbance and Patch Dynamics: An Introduction. In: Pickett STA, White PS, editors. The Ecology of Natural Disturbance and Patch Dynamics. San Diego: Academic Press. p. 3–13.

Yang W, Feiner N, Salvi D, Laakkonen H, Jablonski D, Pinho C, Carretero MA, Sacchi R, Zuffi MAL, Scali S, et al. 2022. Population Genomics of Wall Lizards Reflects the Dynamic History of the Mediterranean Basin. Molecular Biology and Evolution 39.

Yiming L, Siqi W, Chaoyuan C, Jiaqi Z, Supen W, Xianglei H, Xuan L, Xuejiao Y, Xianping L. 2021. Latitudinal gradients in genetic diversity and natural selection at a highly adaptive gene in terrestrial mammals. Ecography 44:206–218.

