## Supplemental Information for "Large scale, low coverage population genomics approach to assess genetic variation and structure across vulnerable Swedish sand lizard populations"

### Table of Contents:


| | |
|---|---|
| **Figure A1:** Coverage profiles | Page 2 |
| **Figure A2:** Genetic load across populations | Page 3 |
| **Figure A3:** Genetic load proportions across latitude | Page 4 |
| **Figure A4:** Admixture plots for K=2 to K=7 | Page 5 |
| **Figure A5:** Admixture plots for K=8 to K=13 | Page 6 |
| **Figure A6:** Admixture delta k | Page 7 |
| **Figure A7:** FST scans | Page 8 |
| **Table A1:** Population information | Page 9 |
| **Table A2:** Global $F_{ST}$ for population pairs | Page 9 |

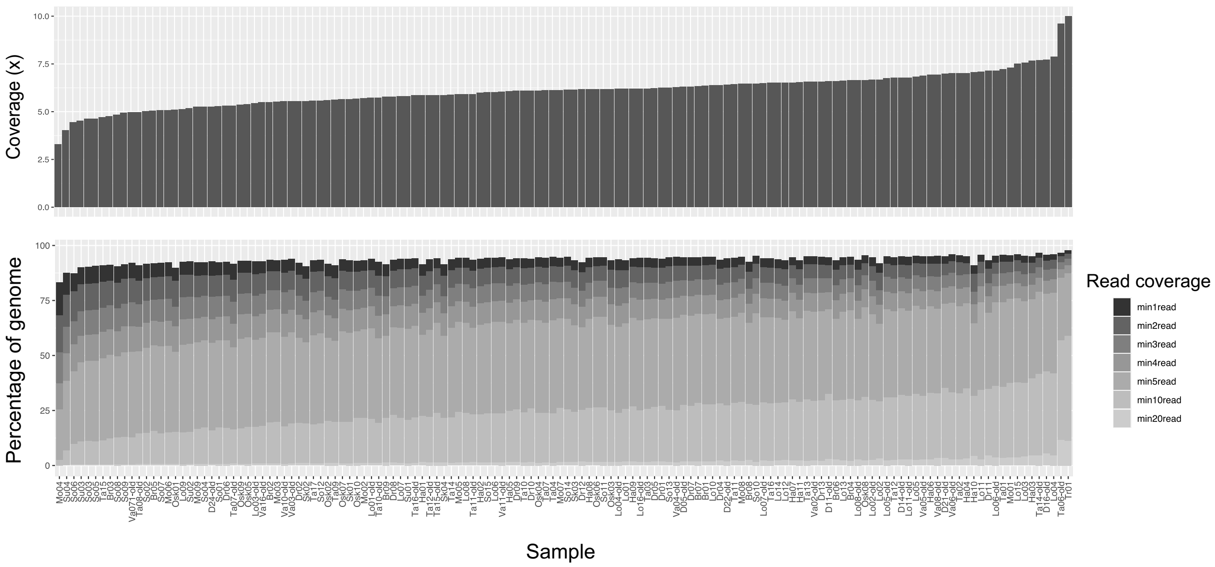


**Figure A1**. Coverage profiles for whole genome sequencing of sand lizard samples. Upper plot shows x coverage per sample. Lower plot displays percentage of genome covered by at least 1, 2, 3, 4, 5, 10 and 20 reads. Both plots are in same sample order of increasing x coverage.

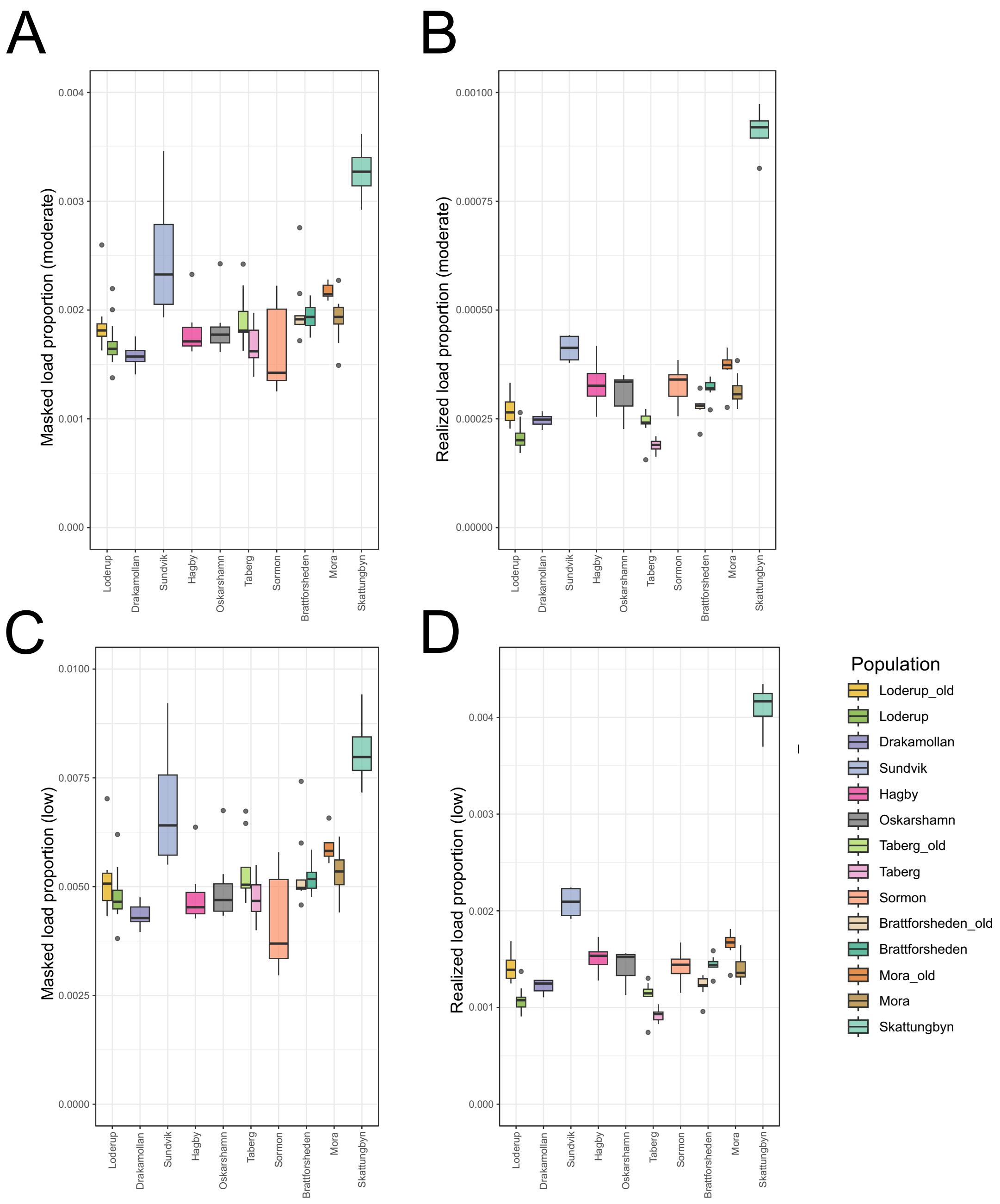


**Figure A2.** Genetic load across Swedish sand lizard populations, including masked genetic load proportions for moderate effects (A), realised genetic load proportions for moderate effects (B), masked genetic load proportions for low effects (C), and realised genetic load proportions for low effects (D). Populations are ordered by increasing latitude.

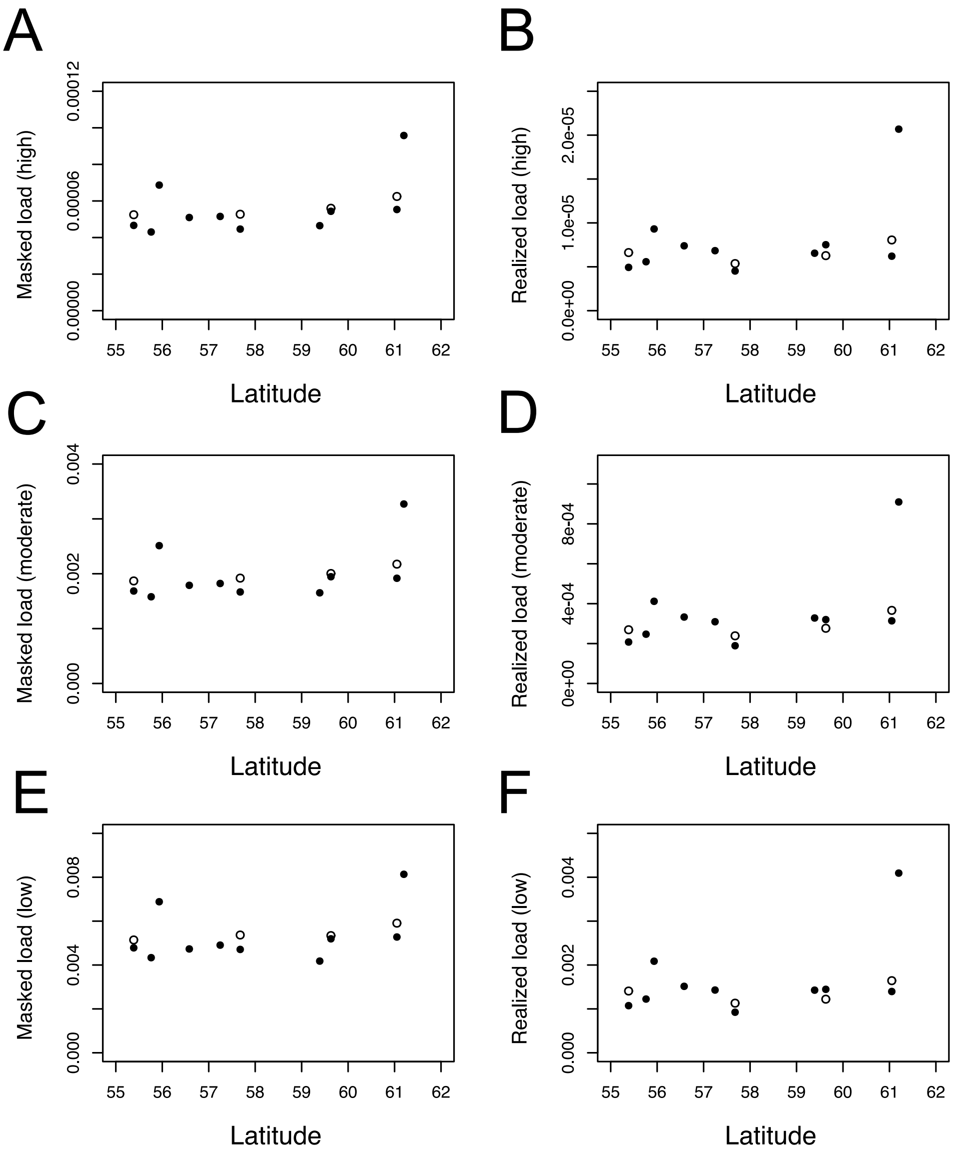


**Figure A3.** Genetic load proportions across latitude, including mean masked genetic load proportion with high impact (**A**), mean realised genetic load proportion with high impact (**B**), mean masked genetic load proportion with moderate impact (**C**), mean realised genetic load proportion with moderate impact (**D**), mean masked genetic load proportion with low impact (**E**), mean masked genetic load proportion with low impact (**F**). Contemporary population samples indicated by filled circles. Earlier sampling indicated by open circles.

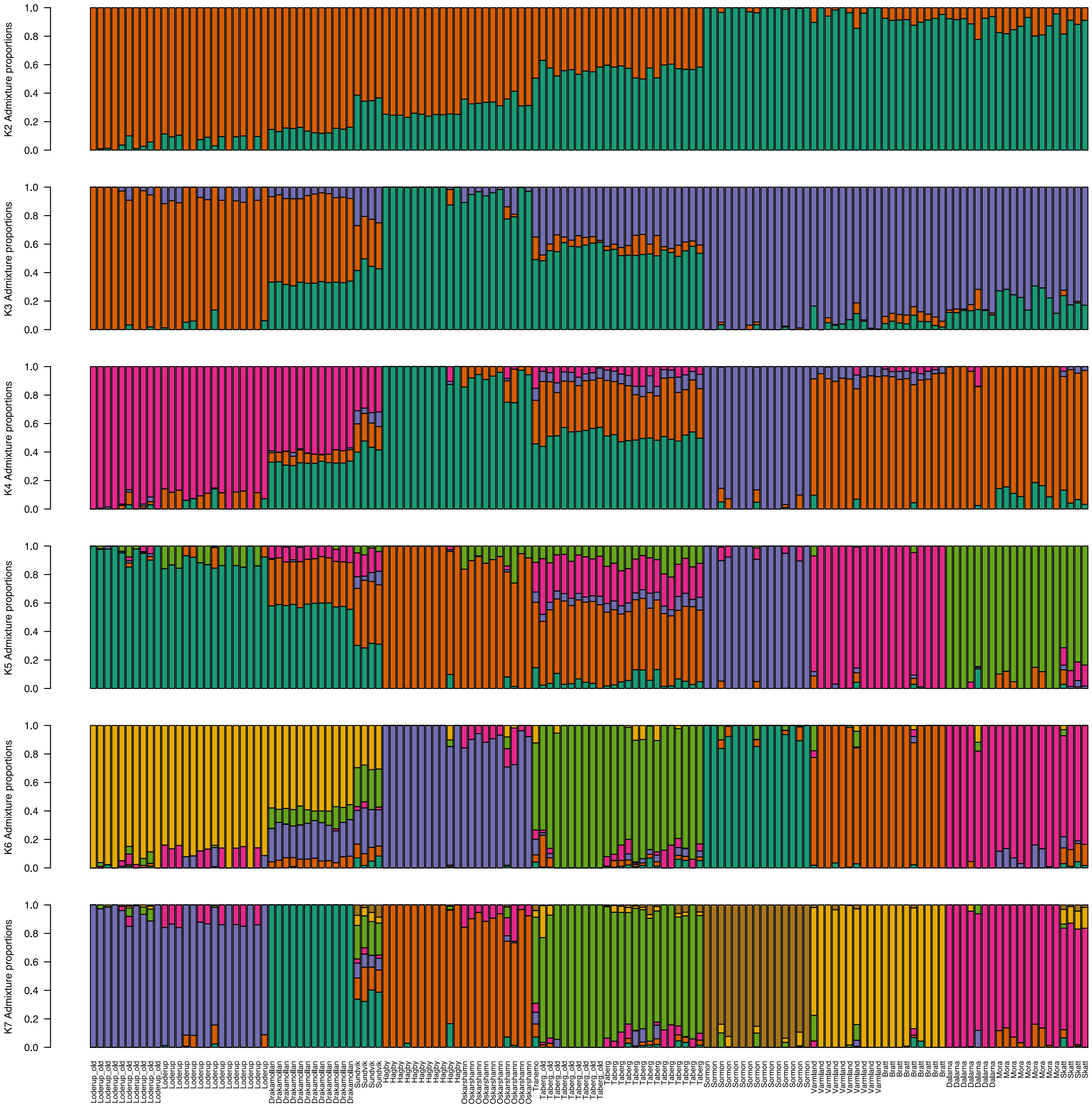


**Figure A4**. Admixture plots from K=2 to K=7. Ancestry proportions of individuals as estimated by NGSadmix software for K=2, K=3, K=4, K=5, K=6 and K=7. Population of origin indicated for each individual below barplots. Populations ordered by latitude.

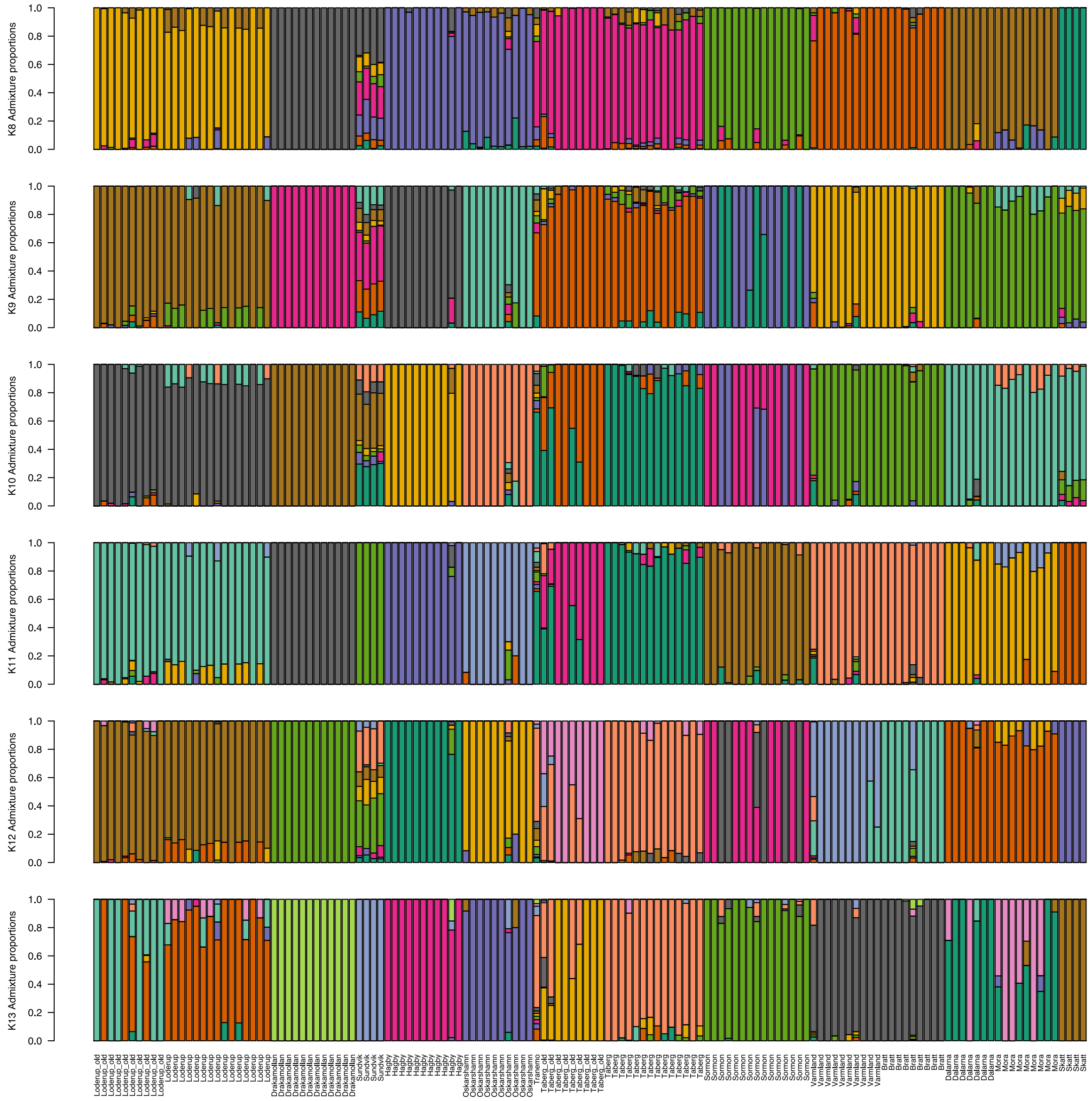


**Figure A5**. Admixture plots from K=8 to K= 13. Ancestry proportions of individuals as estimated by NGSadmix software for K=8, K=9, K=10, K=11, K=12 and K=13. Population of origin indicated for each individual below barplots. Populations ordered by latitude.

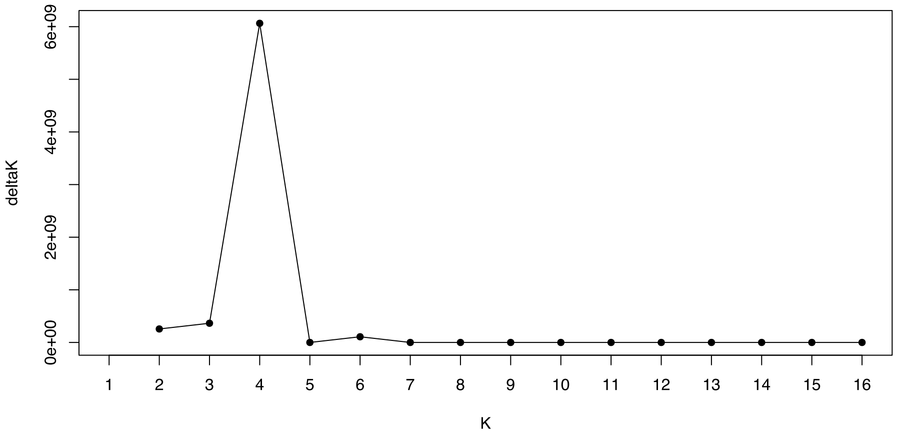


**Figure A6**. Best *K* of 4 estimated from using delta*K* from resulting likelihood values

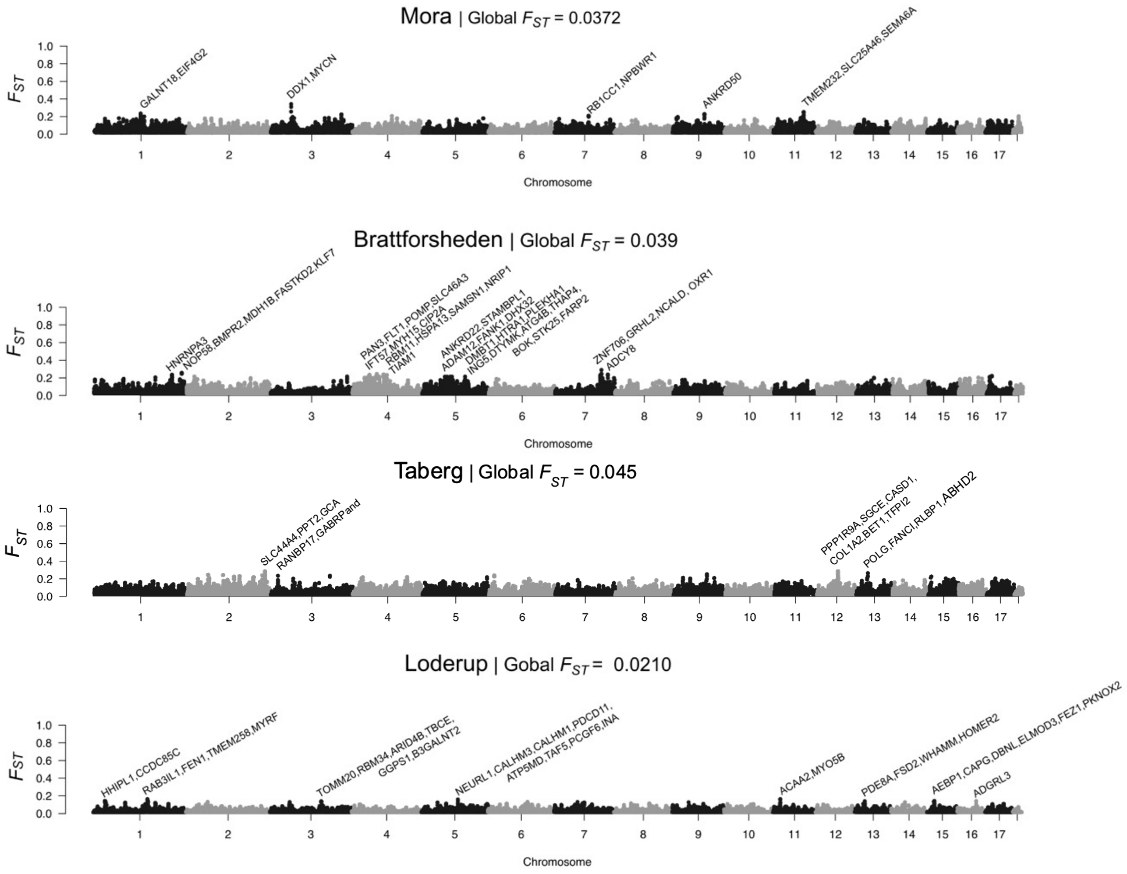


**Figure A7**. $F_{ST}$ scans. Pairwise global $F_{ST}$ estimates and window-based $F_{ST}$ scans (100 kb windows with 50 kb step size) estimated within populations that had been sampled at two separate time points, including (from upper to lower plots) Mora, Brattforsheden, Taberg, Löderup. Global $F_{ST}$ within populations over time was low, allowing for investigation of potential regions under selection. Genes within putative regions under selection (within $F_{ST}$ peaks) with an allele frequency difference greater than 0.6 are indicated in the plots.

**Table A1.** Population information, including Swedish county, latitude and longitude

| County | Population | Latitude | Longtitude |
|---|---|---|---|
| Dalarna | Skattungbyn | 61.20 | 14.90 |
| Dalarna | Mora | 61.05 | 14.45 |
| Värmland | Brattforsheden | 59.63 | 13.97 |
| Värmland | Sörmon | 59.39 | 13.35 |
| Jönköping | Taberg | 57.68 | 14.08 |
| Kalmar | Oskarshamn | 57.24 | 16.34 |
| Kalmar | Hagby | 56.58 | 16.19 |
| Skåne | Sundvik | 55.93 | 12.79 |
| Skåne | Drakamöllan | 55.76 | 14.12 |
| Skåne | Löderup | 55.39 | 14.10 |

**Table A2.** Global $F_{ST}$ (weighed) for population pairs sampled in 2019.

| | Skattungbyn | Mora | Brattforsheden | Sörmon | Taberg | Oskarshamn | Hagby | Löderup | Drakamöllan |
|---|---|---|---|---|---|---|---|---|---|
| Mora | 0.2254 | - | - | - | - | - | - | - | - |
| Brattforsheden | 0.2629 | 0.1611 | - | - | - | - | - | - | - |
| Sörmon | 0.2987 | 0.2099 | 0.2020 | - | - | - | - | - | - |
| Taberg | 0.2088 | 0.1254 | 0.1327 | 0.1731 | - | - | - | - | - |
| Oskarshamn | 0.2674 | 0.1660 | 0.1941 | 0.2400 | 0.1248 | - | - | - | - |
| Hagby | 0.2927 | 0.1985 | 0.2084 | 0.2549 | 0.1351 | 0.1182 | - | - | - |
| Löderup | 0.2657 | 0.1765 | 0.1880 | 0.2369 | 0.1268 | 0.1563 | 0.1625 | - | - |
| Drakamöllan | 0.2668 | 0.1805 | 0.1751 | 0.2333 | 0.1154 | 0.1435 | 0.1478 | 0.0904 | - |
| Sundvik | 0.3097 | 0.2113 | 0.2107 | 0.2563 | 0.1334 | 0.1725 | 0.1869 | 0.1594 | 0.1456 |
